# Neuron Geometry Underlies Universal Network Features in Cortical Microcircuits

**DOI:** 10.1101/656058

**Authors:** Eyal Gal, Rodrigo Perin, Henry Markram, Michael London, Idan Segev

**Affiliations:** The Edmond and Lily Safra Center for Brain Sciences, the Hebrew University of Jerusalem, Israel; Department of Neurobiology, the Hebrew University of Jerusalem, Israel; Blue Brain Project, École Polytechnique Fédérale de Lausanne, Geneva, Switzerland

**Author notes:** Corresponding authors. (E.G.); (I.S.).

## Abstract

Why do cortical microcircuits in a variety of brain regions express similar, highly nonrandom, network motifs? To what extent this structure is innate and how much of it is molded by plasticity and learning processes? To address these questions, we developed a general network science framework to quantify the contribution of neurons’ geometry and their embedding in cortical volume to the emergence of three-neuron network motifs. Applying this framework to a dense *in silico* reconstructed cortical microcircuits showed that the innate asymmetric neuron’s geometry underlies the universally recurring motif architecture. It also predicted the spatial alignment of cells composing the different triplets-motifs. These predictions were directly validated via *in vitro* 12-patch whole-cell recordings (7,309 triplets) from rat somatosensory cortex. We conclude that the local geometry of neurons imposes an innate, already structured, global network architecture, which serves as a skeleton upon which fine-grained structural and functional plasticity processes take place.

## INTRODUCTION

Anatomical connectivity in brain circuits plays a key role in governing network dynamics and emerging brain functions(Avena-Koenigsberger et al., 2017; Bonifazi et al., 2009). Connectomes at all scales exhibit structured, highly nonrandom, connectivity patterns(Bassett and Sporns, 2017; Fulcher and Fornito, 2016; Perin et al., 2011; Rieubland et al., 2014; Song et al., 2005). Specifically, a glimpse into the microscale circuitries in several brain regions was provided recently using *in vitro* whole-cell multi-patch recordings(Perin et al., 2011; Rieubland et al., 2014; Song et al., 2005) (**Fig. 1a**) and by *in silico* dense reconstructions of cortical microcircuits(Egger et al., 2014; Markram et al., 2015) (**Fig. 1b**). Surprisingly, all observed microcircuits display common network architecture, over-expressing specific local connectivity patterns (motifs; **Fig. 1c**). However, despite their cross-region universality, the origin of these motifs has remained elusive.

**Figure 1.**
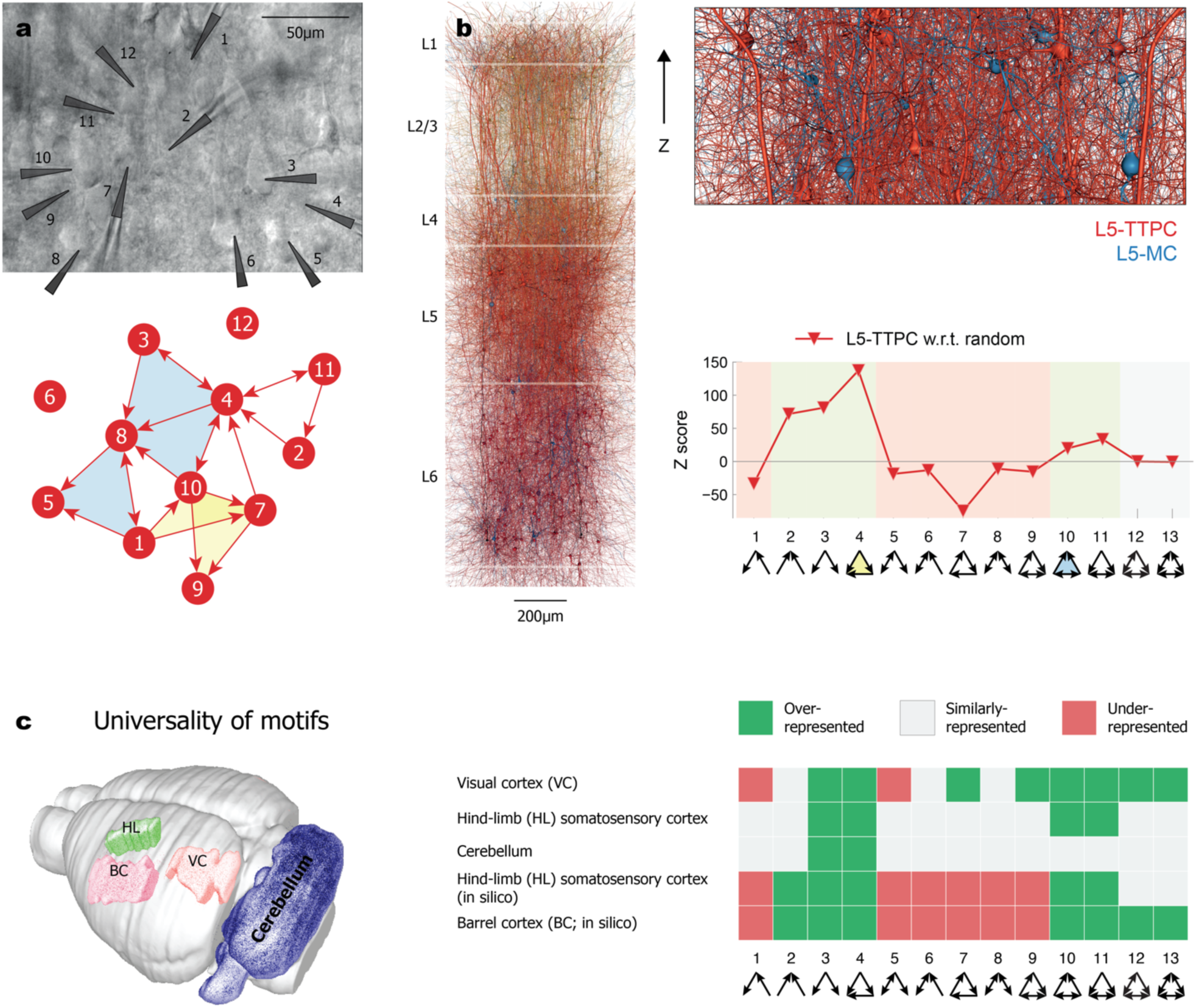
Universality of triplet motifs appearance in a variety of mammalian neuronal microcircuits. **a**, Differential contrast video image of *in vitro* 12-patch whole-cell recording from L5 thick tufted cortical pyramidal cells in rat somatosensory cortex (top). Triggering a spike in a presynaptic cell and recording excitatory post-synaptic synaptic potentials in the post-synaptic cells enabled us to establish circuit connectivity diagram (schematically depicted at the bottom) and to count the frequencies of specific triplet motifs (two types of motifs are highlighted by light green and yellow). **b**, Left - dense *in silico* reconstruction of neocortical microcircuit (NMC) from the rat somatosensory cortex(Markram et al., 2015); its volume is of ∼0.3 mm^3^ with ∼36 million synapses. Top right - a slice in L5 from the NMC circuit showing L5 thick tufted pyramidal cells (L5-TTPC; red) and L5 Martinotti cells (L5-MC; blue). Lower frame – a graph theoretical analysis of the connectivity diagrams for the two cell types shown above, highlighting the overrepresented triplet motifs (e.g., motifs #3 and #4), the underrepresented ones (e.g., motif #7 for L5-TTPCs), and those that are similarly represented, as compared to the respective distance-dependent random network (see **Methods**). Triplet #4 is marked in yellow and triplet #10 in light blue, as in **a. c**, Universality of overrepresented triplet motifs in a variety of neuronal microcircuits. Note that triplets #3 and #4 are overrepresented in all circuits. The brain regions in which these microcircuits reside are depicted schematically on the left; the expression of the various triplet motifs in these circuits is shown on the right. Triplet motif expression data are displayed for primary visual cortex(Song et al., 2005); somatosensory cortex(Perin et al., 2011); interneurons in the cerebellum(Rieubland et al., 2014); dense *in silico* somatosensory cortex(Gal et al., 2017), and for dense *in silico* barrel cortex(Egger et al., 2014).

In the absence of a concrete theory about the principles underlying these structured connectivity patterns in neuronal microcircuits, their emergence was considered as being shaped by specific genetic markers(Yamagata and Sanes, 2008; Yu et al., 2009) or by plasticity and learning processes(Ocker and Doiron, 2018; Ravid Tannenbaum and Burak, 2016; Rieubland et al., 2014; Song and Abbott, 2001). However, these may not be the only mechanisms by which the connectivity patterns emerge. In fact, most types of cortical neurons display highly asymmetric geometry with dendritic and axonal trees typically extending in different directions. This asymmetric morphology may, by itself, introduce anisotropy to the probability for forming pairwise connections between neurons in the circuit, and, consequently, “enforce” the existing complex profile of higher-order cortical motifs. If this is the case, then plasticity and learning processes might operate on top of an innate, pre-structured, cortical skeleton of network architecture rather than on a *tabula rasa* organization. This innate architecture would impose a strong constraint on the extent to which plasticity could further shape network connectivity. Motivated by this parsimonious hypothesis for the universality of network local architecture, in this work we combine theoretical modeling and electrophysiological recording to study the extent to which geometry of neurons alone dictates the ubiquitous neuronal-specific profile of over- and under-represented motifs at the microcircuit level.

To test this challenging possibility theoretically, we utilized a reference circuit of a dense *in silico* reconstruction of a 0.3 mm^3^ volume from rat somatosensory cortex(Markram et al., 2015) (the neocortical microcircuit, NMC). The NMC was used for two main reasons. First, it reproduces a triplet motif profile that is consistent with those found experimentally(Gal et al., 2017). The emergence of these triplets in the NMC, whose connectivity is based on strictly pairwise geometrical rules between 3D-reconstructed neurons, hints of their geometric origin. Second, the unprecedented magnitude of cellular level connectomics data, mapping ∼36 million synapses interconnecting ∼31,000 neurons, enables the identification of regularities in the network architecture and advanced modeling to uncover the underlying organizing principles. Here, through a systematic construction of a series of progressively more complex generative geometrical circuit models, we show that the asymmetrical morphology of neurons and their embedding in the cortical 3D space is indeed sufficient to explain the preference for common motifs in a variety of neuronal microcircuits.

Next, we experimentally studied the appearance of triplet motifs using 12-patch whole-cell recordings from layer 5 thick tufted pyramidal cells (L5-TTPC) in the rat somatosensory cortex. As predicted by our geometrical models, we found in the biological tissue a close match between the physical position of L5-TTPCs triplets and the motif they form. This implies that the local geometry of L5-TTPCs and their distribution in the 3D cortical volume impose an innate global and highly structured skeleton in the L5-TTPCs circuit.

## RESULTS

### Origin of triplet motifs in dense *in silico* cortical microcircuits

What could be the origin for such universality in the prevalence of overrepresented triplet motifs? We examined this question theoretically using the NMC as a benchmark circuit. Connectivity in the NMC is based on the proximity of axons and dendrites (“touch detection” algorithm) together with a set of synaptic pruning rules, which are set to maintain an observed distribution of the number of contacts per connection but are consistent with a purely geometry-driven connectivity pattern(Markram et al., 2015; Reimann et al., 2015). Therefore, the emergence of triplet motifs in this circuit must arise from certain geometrical rules (see **Discussion**). We uncovered these rules by building a series of network models which progressively consider higher-order features of the geometry underlying neuronal connectivity in the NMC, exploring whether these triplet motifs will emerge in these generative network models as they do in the NMC target circuit (**Fig. 2, Fig. S1**).

**Figure 2.**
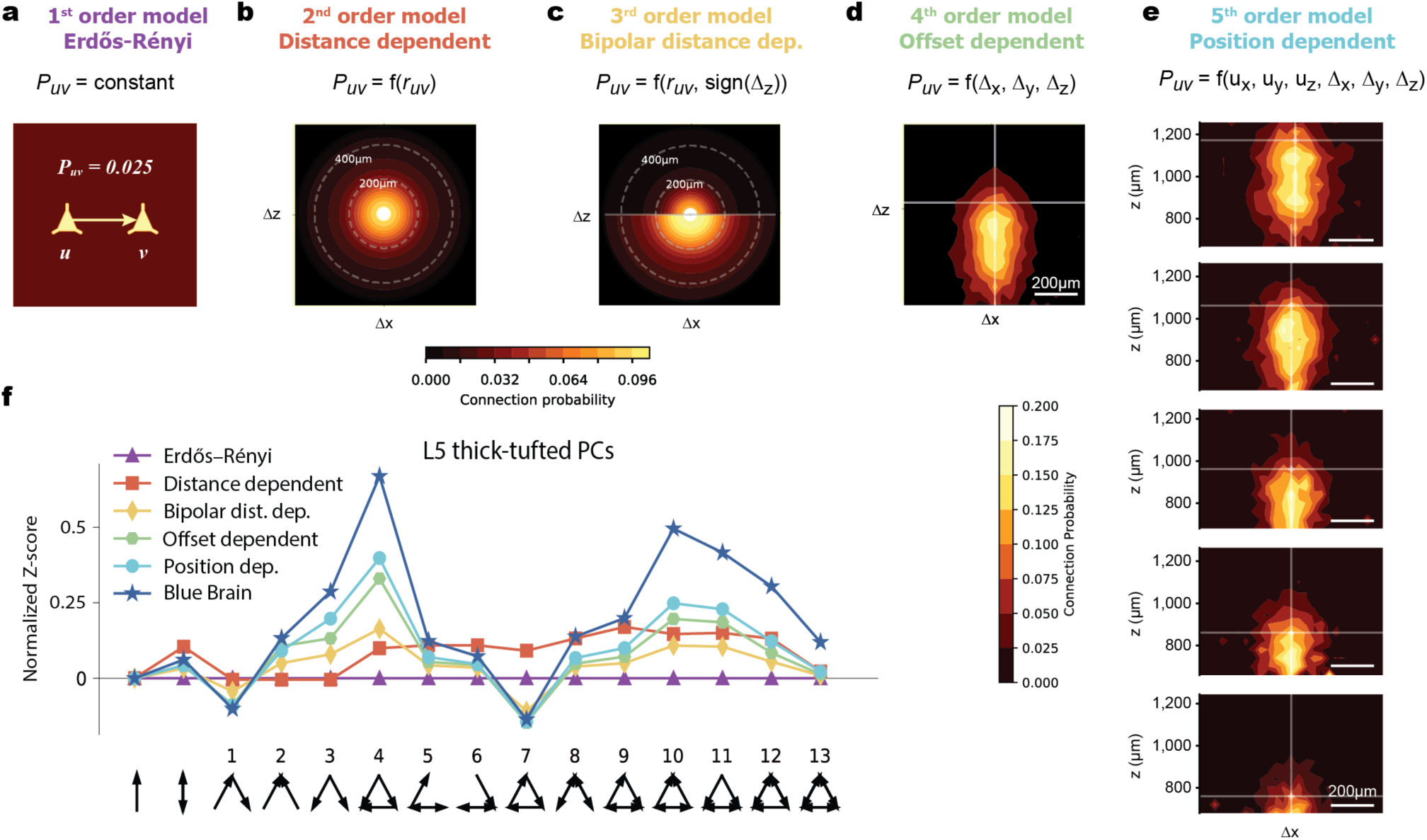
The anisotropic geometry of cortical neurons underlies the emergence of triplet motifs. **a-e**, a series of progressively more complex generative models for the circuit connectivity of L5 thick-tufted pyramidal cells (L5-TTPC). **a**, 1^st^ order Erdős-Rényi model, with uniform connection probability, *p* = 0.025, among the cells as is the average connection probability in the NMC. **b**, 2^nd^ order model, in which *p* (color coded) is distance dependent, as derived from the NMC. *p* is depicted with respect to the distance from the postsynaptic cell located at the origin (Δx, Δz intersection point). Note the decrease in *p* with distance. **c**, 3^rd^ order “bipolar” model, in which *p* also depends on whether the postsynaptic neuron is above or below the presynaptic cell (along the z-axis). Note that, in the NMC, *p* is larger for downwards connection. **d**, 4^th^ order “offset dependent” model, in which *p* depends also on the relative direction, in 3D, to the postsynaptic cell as it does in the NMC. Note that *p* is larger for postsynaptic neurons that are directly below (along the z-axis), as compared to cells that are obliquely below the presynaptic cell. **e**, 5^th^ order model, in which *p* depends on the absolute position in 3D of both the presynaptic and the postsynaptic cells. E.g., L5-TTPCs whose somata are located close to the border of layer 4 (top horizontal line) have a larger span, in the z-direction, for having a postsynaptic partner as compared to L5-TTPCs whose somata reside near the border of layer 6 (bottom). **f**, Distribution of the various triplet motifs for each of the models shown in a-e, with respect to the reference Erdős-Rényi model (x-axis). Motif distribution in the NMC is shown by the blue line. Note that, as the geometry captured by the model becomes more realistic, the emerging motif profile becomes closer to that of the NMC (blue line with stars). See Methods for a detailed explanation regarding the construction of the different generative models used in this figure.

We started by focusing our attention on L5-TTPCs and modeled their interconnectivity by the simplest, 1^st^ order, random network: an Erdős-Rényi random network preserving same average connection probability of p = 0.025 as in the NMC (**Fig. 2a**, see also (Egger et al., 2014; Gal et al., 2017; Perin et al., 2011; Rieubland et al., 2014; Song et al., 2005)). This random model was not sufficient to capture the over- and under-represented triplet motifs empirically observed in the NMC (P < 0.001, Monte Carlo; **Fig. 2f**, purple vs. blue line).

Next, we constructed a 2^nd^-order random network which, in addition to the average connection probability, also preserves the dependence of connection probability on intersomatic distance as in the NMC (**Fig. 2b**, see also (Gal et al., 2017; Perin et al., 2011; Rieubland et al., 2014)). Specifically, the probability for a connection from a neuron (*u*) to another (*v*) obeys

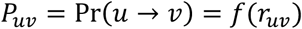

Where *r*_*uv*_ is the intersomatic distance between neurons *u* and *v*. This extended distance-dependent random model also does not explain the profile of motifs that were found in the NMC (P < 0.001, Monte Carlo; **Fig. 2f**, red line). In particular, the feedback loop motif (#7), which is underrepresented in the NMC, is incorrectly overrepresented in the 2^nd^-order random network. It is important to note that, in both Erdős-Rényi and distance-dependent models, the probability for the feedback loop motif is proportional to the probability for the feedforward motif (#4) by definition. Hence, in both models, it is impossible to decrease the prevalence of motif #7 without decreasing motif #4 too. Thus, to better fit the empirical motif profile, a new model would have to break the symmetry between these two motifs.

The next 3^rd^-order model also considers the implication of the asymmetrical geometry of L5-TTPCs whose dendrites tend to project upwards towards the pia (in the z-direction) and their axons that tend to project in the opposite direction. Because the prerequisite for the formation of a synapse is that the axon of the presynaptic cell should come close to the dendrite of the post-synaptic cell, the probability of connection between pairs of L5-TTPCs in the NMC is larger in the downwards (z) direction. The simplest way to allow such bias in the probability function is by introducing a dependency on an additional variable indicating whether the target cell is above or below the source cell (sign(Δ_*z*_)). Thus, the probability for a connection takes the form

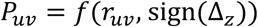

Indeed, taking into account this bipolar effect of dendrites and axons when quantifying the connection probability in a 3^rd^-order model uncovers a highly anisotropic connectivity function (**Fig. 2c**). This model starts to capture the motifs trends as is the case of the NMC. Specifically, motifs #3, #4, #5, and #6 are overrepresented with respect to the Erdős-Rényi reference model, whereas motif #7 is correctly underrepresented (**Fig. 2f**, yellow line). Thus, by taking into consideration the anisotropy imposed by the bipolar nature of L5-TTPCs, the symmetry between motifs #4 and #7 is properly broken.

In our 4^th^-order model, we further added the dependency of connection probability on the complete 3D offset between the two cells (**Figure 2d** demonstrated the offset in an x-z cross section)

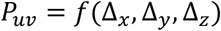

This further improved the similarity of motif appearances between the model and the NMC (**Fig. 2f**, green line). To ensure that the improvement indeed results from extra information captured by the geometric offset features, and not merely as a consequence of a larger number of features, the connection probability functions of all geometrical models were fitted using a machine learning classifier having the same complexity (**Methods**).

Finally, in our 5^th^-order model, we additionally considered the absolute positions of the cells when predicting connections

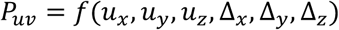

L5-TTPCs whose somata are located at the upper part of layer 5 and have a more elongated probability function, in the z-direction, than deeper cells (**Fig. 2e**). When this effect was incorporated, the reproduced motif profile was even closer to that of the NMC (*P* <0.001, two-tailed Wilcoxon rank sum test; **Fig. 2f**, light blue line).

### Geometrical considerations explain the emergence of cortical motifs

The similarity between triplet distribution in experiments and the *in silico*, geometrically based, NMC model, strongly suggests that the emergence and frequency of triplet motifs in the biological tissue can be explained, for the most part, as solely due to geometrical considerations. The emergence of triplet motifs for highly polar neurons, such as L5-TTPCs (**Fig. 3a**) and L5 Martinotti cells (L5-MCs) (**Fig. 3c**), depends strongly on their exact bipolar structure and position in 3D. When the neuron is more isotropic, as is the case for the inverted L6 pyramidal neurons (in which dendrites and axons tend to overlap), then the anisotropic models (beyond the 2^nd^-order distance-dependent model) do not provide further improvement for explaining the triplet distribution of networks composed of such cells (**Fig. 3b,d**).

**Figure 3.**
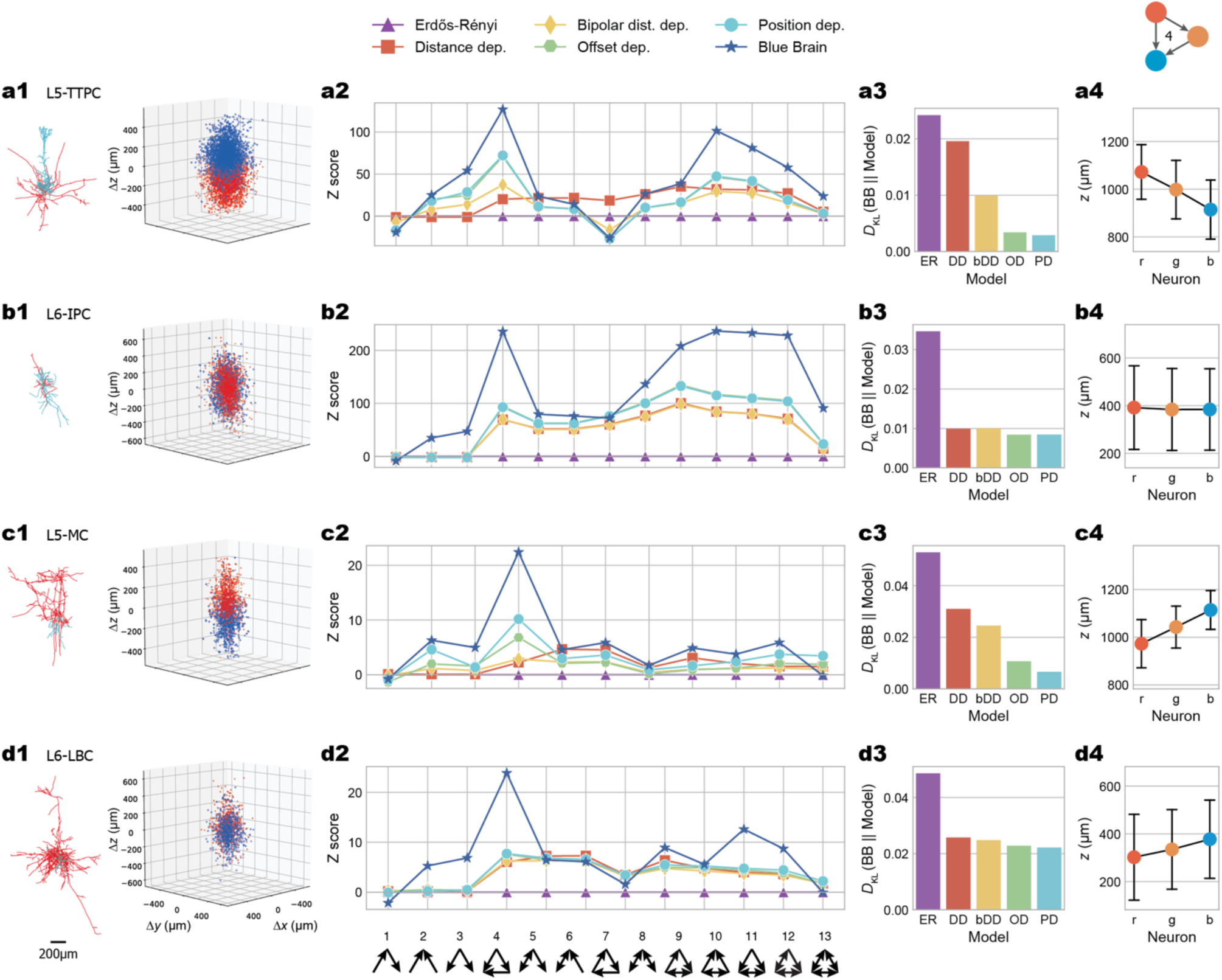
Impact of cell-type-specific morphology on the triplet motif profile. **a**, Appearance of triplet motifs for L5-TTPC. **a1**, Left, an exemplar 3D reconstructed TTPC cell. Right, the relative soma position of all the presynaptic TTPC cells (blue) and all the postsynaptic TTPC cells (red) with respect to the cell somata centered at 0,0 (superposition of 50 TTPC cells). **a2**, Distribution of triplet motifs for TTPC cells (n=2,003). **a3**, Kullback–Leibler divergence between the motif distribution observed in the NMC and each of the 5 models shown in A2. Note the gradual improvement with model complexity. **a4**, spatial embedding of neurons composing motif #4 (mean ± s.d.). **b-d**, As in **a**, but for **b**, L6 inverted pyramidal cells (n = 3,476), **c**, L5 Martinotti cells (n=395) and **d**, L6 large basket cells (n=463). Note that when the neuron is more isotropic as in **b** and **d**, models beyond the distance-dependent model do not provide further improvement.

We thus concluded that the skeleton of circuit motifs is dictated, innately, by geometrical considerations; namely, by the position of the neurons in 3D and the asymmetry of their morphology. This implies that processes such as plasticity and specificity of cortical connections play a second order role in shaping motif structure and frequency in these circuits.

### Experimental validation of motifs spatial organization in L5 cortical pyramidal cells

Interestingly, the geometrical models imply a specific spatial organization for each motif. For instance, the anisotropic models predict that, for vertically polarized neurons, the feedforward motif (#4) will be vertically oriented in space (**Fig. S2)**. Specifically, for L5-TTPCs the “source” neuron (the presynaptic neuron that projects to the other two neurons) is predicted to be located above the other two cells whereas the “sink” neuron (the postsynaptic neuron receiving synapses from the other two neurons) predicted to be the lowest (**Fig. S2a**). Indeed, we found a tendency for such an alignment in the NMC circuitry (**Fig. 3a4**). In addition, for the L5-MCs, whose vertical polarization is inverted (axons stretch mostly above the soma whereas the dendrites stretch mostly below it) the models predict an inverted vertical directionality of neuron locations (**Fig. 3c4, Fig. S2b**).

To examine experimentally our predictions about the spatial embedding of motifs we recorded *in vitro* from rat somatosensory cortex L5-TTPCs and reconstructed both the synaptic connectivity among the neurons and their spatial 3D position. (**Fig. 4a**). Parasagittal slices oriented parallel to the apical dendrites were used in order to maintain morphologies maximally intact. Using whole-cell recordings of up to 12 cells patched simultaneously we mapped and quantified the connectivity in 7,309 triplets (an example is shown in **Fig. 4, a-c)** by stimulating with trains of pulses each one of the patched cells in turn and averaging the postsynaptic cell responses to 15-30 rounds of stimulation. Repeated excitatory postsynaptic responses with latencies of 1-3ms and amplitudes above 100μV were used as indicators of monosynaptic connections (**Methods**).

**Figure 4.**
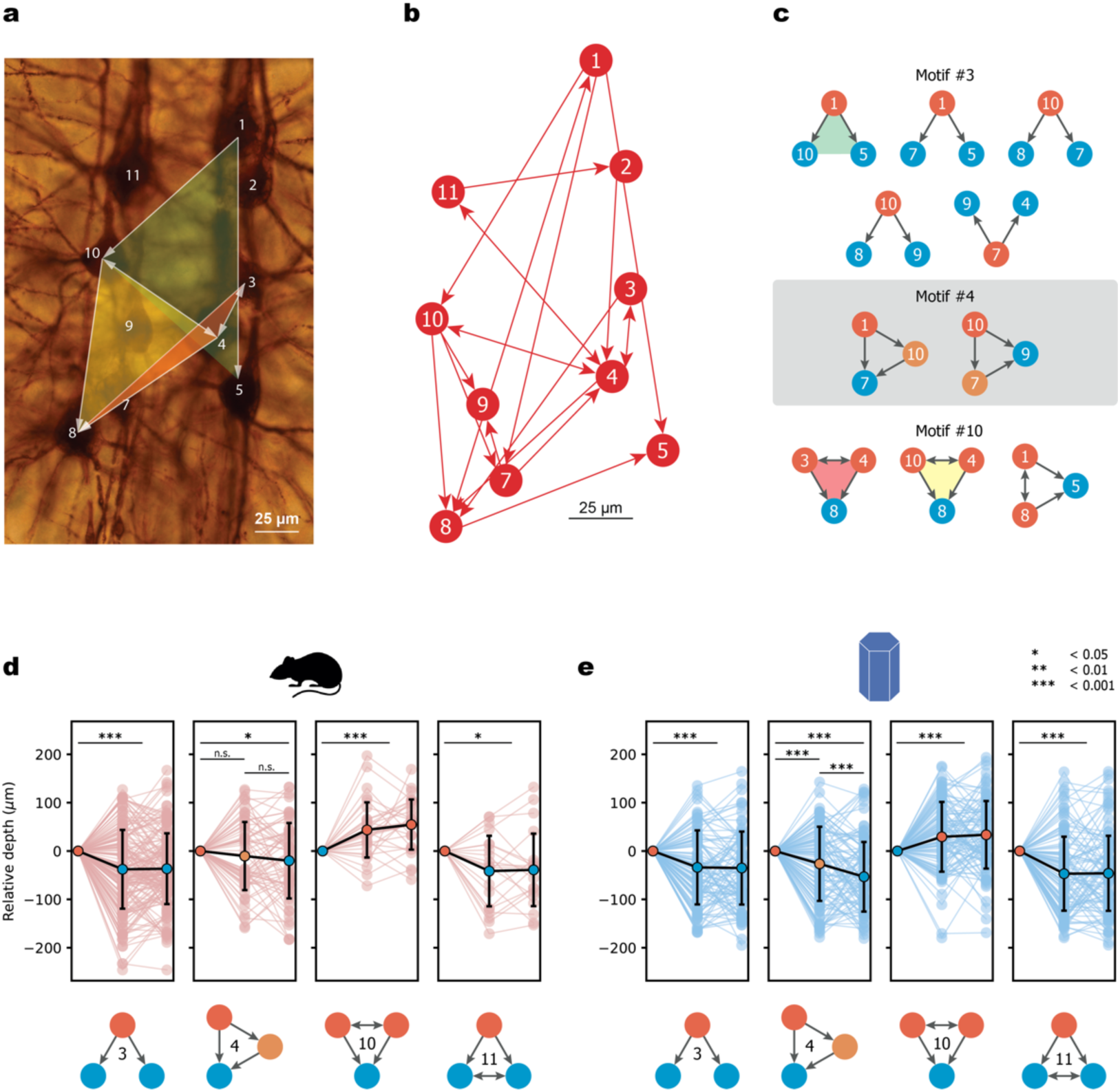
Experimental validation of motif spatial embedding. **a**, Biocytin staining of multi-patch recordings from 12 L5-TTPCs. Three exemplar connected triplet cells are shown by the transparent (green, yellow and red) triangles. **b**, Connectivity map for the 12-neuron circuit shown in **a. c**, All instances for three triplet motifs (#3, #4 and #10) that were extracted from the circuit diagram shown in **b**. The relative depth (z-axis) of the cells composing these triplets is preserved. E.g., note that, for triplet #3, in most (but not in all) cases the presynaptic cell (red circle) is located above the two postsynaptic cells (blue circles). Green, yellow and pink triangles correspond to the respective cell triplets shown in **a. d**, Spatial embedding of triplet motifs (#3, #4, #10 and #11) for L5-TTPCs in the experimental data set (*n*_3_ = 163, *n*_4_ = 63, *n*_10_ = 31, *n*_11_ = 21 triplets); the relative depth of the cells forming individual triplets (light brown lines) and mean ± s.d. (black line). *\*P* < 0.05, \*\**P* < 0.01, \*\*\**P* < 0.001 (two-sided paired-sample t-test). P values for R-vs-Y and Y-vs-B in motif #4 are 0.2470 and 0.3029, respectively. For clarity, cell depths were aligned in each triplet according to one member (the left-most cell in the plot). **e**, As in **d** but for a 400μm-thick slice from the NMC (*n*_3_ = 219,377, *n*_4_ = 22,137, *n*_10_ = 833 and *n*_11_ = 765 triplets of which only 100 instances are shown). Note the very close similarity between experiments (**e**) and model (**d**).

The *in vitro* data indeed shows that, as predicted by our geometrical models, there is a close match between the physical position of L5-TTPCs triplets in space and the synaptic connectivity motif they form in the biological tissue (**Fig. 4d)**. A strikingly similar tendency for the spatial embedding of motifs was also observed among L5-TTPCs in the NMC (**Fig. 4e, Methods**). The close similarity in the average relative depth differences in the experimental and NMC triplet patterns lends support to the notion that neuronal geometry is a fundamental factor influencing/determining network architecture.

## DISCUSSION

Neuronal circuits in a variety of mammalian brain areas share a similar highly nonrandom expression pattern of network motifs (**Fig. 1**). This observed connectivity structure could be innate and/or the consequence of active plasticity and learning processes. Our study shows that the universal tendency of networks of pyramidal neurons for overrepresenting transitive triplet motifs while underrepresenting intransitive triplets emerges from the asymmetric geometry of the morphology of neurons and their relative spatial positions (**Fig. 2**). The impact of different neuron geometries on the emerging motifs was further evaluated using *is silico* circuits of several other cortical cell types (**Fig. 3**). Theoretical predictions for the spatial alignment of cells were then directly validated via *in vitro* simultaneous whole-cell recordings from up to 12 neurons in the rat somatosensory cortex (**Fig. 4**). We therefore concluded that the spatial organization of neurons and their asymmetric geometry entails most of the motif architecture found in neuronal circuits.

The present study demonstrates that, given the innate bipolar morphology of neurons(Barnes and Polleux, 2009), a structured network connectivity is essentially unavoidable and the observed complex profile of over/underrepresented motifs in brain microcircuitry is largely innate. Additional possible sources of the complex motif profile are structural synaptic plasticity and activity-dependent modifications to the wiring due to learning. However, while these active processes provide a huge number of potential connectivity patterns, we show here that geometry puts strong constraints on the connectivity configuration that are likely, and places a strong bias for the formation of certain types of motifs. Hence, and in agreement with the close similarity that was found between the model predictions and the *in vitro* results, while plasticity and learning may change the specific members of a triplet motif, they are likely to keep the global pattern of motif expression that was set by geometry.

The fine structure of synaptic connections among neurons has been shown to affect neuron-level computations(Amsalem et al., 2016; Cossell et al., 2015), network-level dynamics and information processing(Denk et al., 2012; Zhao et al., 2011), and, ultimately, behavior(Bassett and Sporns, 2017; Seung, 2009). Therefore, it is essential for future studies to consider the contribution of the inevitably overrepresented network motifs to the network activity dynamics in order to advance the unraveling of the long-standing structure-to-function problem in neuroscience.

The generalization of our motif spatial embedding prediction is also supported by a recent study of the connectivity among cerebellar asymmetrical interneurons(Rieubland et al., 2014). In addition, the preference for local transitive patterns has already been reported in different real-world networks from diverse disciplines(Azulay et al., 2016; Benson et al., 2016; Milo et al., 2002, 2004), including the *C. elegans* connectome, social networks, and the World Wide Web. It would be interesting to study, using the present method, the extent to which this transitivity emerges simply from the underlying geometry of such real-world networks.

Our theoretical analysis is based on a detailed dense *in silico* model of a neocortical microcircuit. This provided us with the opportunity to dissect systematically which geometrical features (asymmetric neuronal morphology, neuron position in the cortical tissue) explain the emergence of patterns in the triplet distribution (**Fig. 2**). By adding increasingly more elaborate aspects to the reference generative theoretical network, we have reproduced the general trends of neuronal motifs as found in the full circuit. Despite obtaining a substantial reduction in the discrepancy between the real and expected profile of motif expression, we were unable to completely predict the absolute numbers of the various motifs as found in the full circuit (**Fig. 2f, Fig. 3, a2-d2**). This residual discrepancy likely stems from the fact that the present models define the probability function based on average values in different subpopulations and neglect instance-level differences among individual neurons(Nolte et al., 2019).

Recent studies of macroscale connectomes have also demonstrated that wiring architecture among brain regions is mostly explained by geometric factors, mainly distance(Betzel et al., 2016; Goulas et al., 2019). At the cellular level, previous works have hypothesized that connectivity probably depends on both the distance and the relative orientation between cells(Itzkovitz and Alon, 2005; Rieubland et al., 2014; Schroter et al., 2017); the present study systematically quantifies this dependence and its impact on the topology of the emerging networks and their embedding in cortical space. Additional studies highlight wiring specificities beyond what is expected from simple geometry. Synaptic connections might have a stronger preference to specific cell-type populations(Pfeffer et al., 2013), layers(Harris and Mrsic-Flogel, 2013) or perhaps even to specific cell identity(Kasthuri et al., 2015).

Our theoretical and experimental work provides the foundation for exploring the yet open question regarding the origin of the connectivity structure of cortical microcircuits: what is innate and what is learned? The present work, combined with the soon-coming dense reconstructions of ∼1 mm^3^ of rodents and human cortical microcircuits, will enable us to more fully explore this intriguing question.

## ACKNOWLEDGEMENTS

We thank Oren Amsalem, Toviah Moldwin, Hanan Shteingart and all lab members of the Segev and London labs for many fruitful discussions and valuable feedback regarding this work.

## Funding

This work was supported by the Drahi family foundation to IS, a grant from the EU Horizon 2020 program (720270, Human Brain Project) and a grant from the Gatsby Charitable Foundation. ML is a Sachs Family Lecturer in Brain Science and is supported by the ISF (1024/17) and the Einstein Foundation.

## Author contributions

E.G., M.L. and I.S. conceived the study and wrote the manuscript. E.G. carried out the theoretical analysis. R.P. designed and performed the experiments. H.M. developed the *in silico* microcircuit and provided the respective data.

## Competing financial interests

Authors declare no competing financial interests.

## Data and materials availability

The developed toolbox, together with both the *in vitro* and the *in silico* connectivity data, is available as an open source library at GitHub (https://github.com/gialdetti/netsci). The complete digital reconstruction of the six-layered cortical volume is available at the Neocortical Microcircuit Collaboration Portal (https://bbp.epfl.ch/nmc-portal).

## MATERIALS AND METHODS

### Slice preparation

Experiments were carried out according to the Swiss national and institutional guidelines. Fourteen to eighteen-day-old non-anesthetized Wistar rats were quickly decapitated, and their brains were carefully removed and placed in iced artificial cerebrospinal fluid (ACSF). Slices (300 μm) were cut on an HR2 vibratome (Sigmann Elektronik). Parasagittal slices, ∼1.7–2.2 mm lateral to the midline, were cut to access primary somatosensory cortex (SSC; above the anterior extremity of the hippocampus ± 1 mm). Slices were incubated at 37 °C for 30– 60 min and then left at room temperature until recording. Cells were visualized by infrared differential interference contrast video microscopy using a VX55 camera (Till Photonics) mounted on an upright BX51WI microscope (Olympus). Thick-tufted layer 5 Pyramidal Cells (PCs) were selected according to their large soma size (15–25 μm) and their apparent large trunk of the apical dendrite. Care was taken to use only parallel slices (i.e., slices that had a cutting plane parallel to the course of the apical dendrites and the primary axonal trunk). This ensured sufficient preservation of both the PCs’ axonal and dendritic arborizations.

### Chemicals and solutions

Slices were continuously superfused with ACSF containing 125 mM NaCl, 25 mM NaHCO3, 2.5 mM KCl, 1.25 mM NaH2PO4, 2 mM CaCl2, 1 mM MgCl2, and 25 mM D-glucose bubbled with 95% O2 and 5% CO2. The intracellular pipette solution contained 110 mM potassium gluconate, 10 mM KCl, 4 mM ATP-Mg, 10 mM phosphocreatine, 0.3 mM GTP, 10 Hepes, and 13mMbiocytin adjusted to pH 7.3–7.4 with 5MKOH. Osmolarity was adjusted to 290–300 mOsm L^-1^ with D-mannitol (25–35 mM). The membrane potential values given were not corrected for the liquid junction potential, which is approximately -14 mV. Chemicals were purchased from Sigma Aldrich or Merck.

### Electrophysiological recordings

Multiple somatic whole-cell recordings (4–12 cells simultaneously) were performed with Multiclamp 700B amplifiers (Molecular Devices) in the current clamp mode at 34 ± 1 °C bath temperature. Data acquisition was performed through an ITC-1600 board (Instrutech) connected to a PC running a custom-written routine (PulseQ) under IGOR Pro (Wavemetrics). Sampling rates were 10 kHz, and the voltage signal was filtered with a 2-kHz Bessel filter. Patch pipettes were pulled with a Flaming/Brown micropipette puller P-97 (Sutter Instruments) or a DMZ puller (Zeitz Instruments) and had an initial resistance of 3–8 MΩ.

### Stimulation protocols

Monosynaptic, direct excitatory connections were identified by stimulation of a presynaptic cell with a 20–70 Hz train of 5–15 strong and brief current pulses (1–2 nA, 2–4 ms) followed by a recovery test response 0.5 s after the end of the train (not shown in the traces), all precisely and reliably eliciting action potentials (APs).

### Final somatic positions

The soma positions were recorded relative to an arbitrary reference point, and the z axis was oriented perpendicular to the surface of the slice. After morphological stainings were ready, the y axis was rotated around the z axis to match the orientation of the apical dendrites. The x axis was rotated by the same amount and remained orthogonal to the other two axes.

### Statistics

The statistical significance of the differences between the triplet profile of the NMC and the different generative models was assessed using the Monte Carlo method. Therefore, there were no assumptions made about the data distribution. For each model we estimated the expected motif profile **E** by generating *N* random networks and averaging their emerging motif profiles **e**_*i*_. We then computed the Kullback–Leibler divergence between the NMC observed motif profile **O** and the profile expected by the model E. Monte Carlo *P* values were calculated as (*r* + 1)/(*N* + 1), where *r* is the number of random networks that produced a Kullback–Leibler divergence greater than or equal to that calculated for the NMC (a one-tailed test). We used *N* = 1,000 for each geometrical model.

### Constructing the series of geometrical network models

The observed (neuronal) network of *n* nodes (neurons) is described by *n* × *n* adjacency matrix *A*, whereas the *A*_*ij*_ entry is a binary variable denoting the existence of connection from the neuron *i* to the neuron *i*, and by a 3D vector **x**_*i*_ for each neuron *i*, denoting its 3D position is space, **x**_*i*_ = (*x*_*i*_, *y*_*i*_, *z*_*i*_). Using this data, the geometrical profile of pairwise connection probability function *p* = Pr(*A*_*ij*_ = 1 | **x**_*i*_, **x**_*j*_) was computed at different orders of approximation, gradually taking into account additional features of pair spatial embedding (**x**_*i*_, **x**_*j*_). Then, to study the statistical dependence of network motifs on the geometrical profile a series of generative models for spatially embedded networks was developed, in which each model gradually fits a higher order of approximation of the probability function. The distributions of motif counts emerging from each of the models were compared to the distribution observed in the original network using Kullback–Leibler divergence.

The 1^st^-order model that was used for the generation of reference random networks was the Erdős-Rényi model(Erdös and Rényi, 1959). This model assumes that the probability of forming a connection between two neurons, *p*, is uniform among all pairs of nodes and independent from every other connection. Naïvely, this 1^st^-order model does not take into account spatial features of the nodes whatsoever. Thus, the model requires matching only two parameters: the number of neurons as in the original network, *n* = *n*_*L*5*−TTPC*_ and the mean connection probability over the whole network, *p*(**x**_*i*_, **x**_*j*_) = const = Σ_*i*≠*j*_ *A*_*ij*_ /(*n*(*n* – 1)).

The 2^nd^-order model generated random networks which preserved the non-uniform dependency of connection probability on intersomatic distance (*r*) as observed in the original network (**Fig. 2b, Fig. S1a**). First, the original distance-dependent connectivity profile, *p*^*L*5*−TTPC*^(*r*) (**Fig. S1a**, blue line), was fitted (using an ensemble of 500 decision trees each with a maximal depth of 5 levels (Pedregosa et al., 2011); **Fig. S1a**, orange line). Next, using the fitted probability function, *p*(*r*), we generated random networks with a matching distance-dependent connectivity. The original coordinates of all neurons were preserved, and the connections were defined as independent binomial random variables *A*_*ij*_, with a success probability *p*(**x**_*i*_, **x**_*j*_) = *p*(*r*_*ij*_) that corresponded to their inter-somatic distance *r*_*ij*_.

The 3^rd^-order model captures the fact that the original probability function is anisotropic and depends not only on the intersomatic distance but also on the direction. Specifically, the probability may be different depending on whether the postsynaptic neuron is above or below (along the z-axis) the presynaptic cell. For example, in the L5-TTPC network, the probability is higher when the post-synaptic cell is located below (**Fig. 2c**). The 3^rd^-order model generated networks with a matching “bipolar” distance dependence (**Fig. S1b**).

The 4^th^-order “offset dependent” model captures *p*(**x**_*i*_, **x**_*j*_) dependence on the full 3D relative direction between the presynaptic and postsynaptic cell. Specifically, in the L5-TTPC network, *p*(**x**_*i*_, **x**_*j*_) is larger for postsynaptic neurons that are directly below (along the z-axis), as compared to cells that are obliquely below the presynaptic cell (**Fig. 2d)**.

The 5^th^-order “position dependent” model captures *p*(**x**_*i*_, **x**_*j*_) dependence on the absolute position in 3D of both the presynaptic and the postsynaptic cells. Specifically, L5-TTPCs whose somata are located close to the border of layer 4 (**Fig. 2e**, top horizontal line) have a larger vertical (z-axis) span for connecting a postsynaptic partner as compared to L5-TTPCs whose somata reside near the border of layer 6 (**Fig. 2e**, lower left “cloud”).

In all four (2^nd^-5^th^) geometrical models we employed a machine learning approach to learn the connectivity observed in the NMC. First, given an *n* × *n* adjacency matrix *A* from the NMC, it was transformed into a labelled dataset consisting of *n*(*n* – 1) samples corresponding to all possible pairs of neurons. Each sample is composed of a set of features describing the geometrical embedding of the respective pair (namely, their intersomatic distance, downward/upward alignment, 3D relative offset and absolute 3D positions) and a label indicating whether the pair is connected or not. Secondly, we chose the relevant subset of features for each geometrical model (e.g., for the 3^rd^ order model we chose the intersomatic distance and the downward/upward alignment as the two relevant features). Thirdly, we fitted the probability for a connection from the chosen labelled dataset. Importantly, all 2^nd^-5^th^ order geometrical models were each fitted using an ensemble of 500 decision trees, each with a maximal depth of 5 levels (regression problem)(Pedregosa et al., 2011). This method ensures that the gradual improvement of the fit between the NMC and the progressively more complex geometrical model is indeed a consequence of the additional geometric features and not merely due to the higher degrees of freedom when considering a larger number of features. Finally, using the fitted probability functions, we generated the 2^nd^ – 5^th^ order random networks with matching connectivity as measured in the NMC.

### Spatial embedding analysis

For each motif pattern, we tested the difference between the positions of the composing cells. Specifically, for a given motif, we classified each node according to the number and directions of its connections (different classes are represented by different colors in **Fig. 4, d-e**). Then we performed two-sided paired-sample t-tests between the vertical positions of all different classes. When a motif had two neurons belonging to the same class (e.g., two blue neurons in motif #3) the mean of their vertical position was taken for the purpose of the two-sided paired-sample t-test.

## SUPPLEMENTARY FIGURES

**Figure S1.**
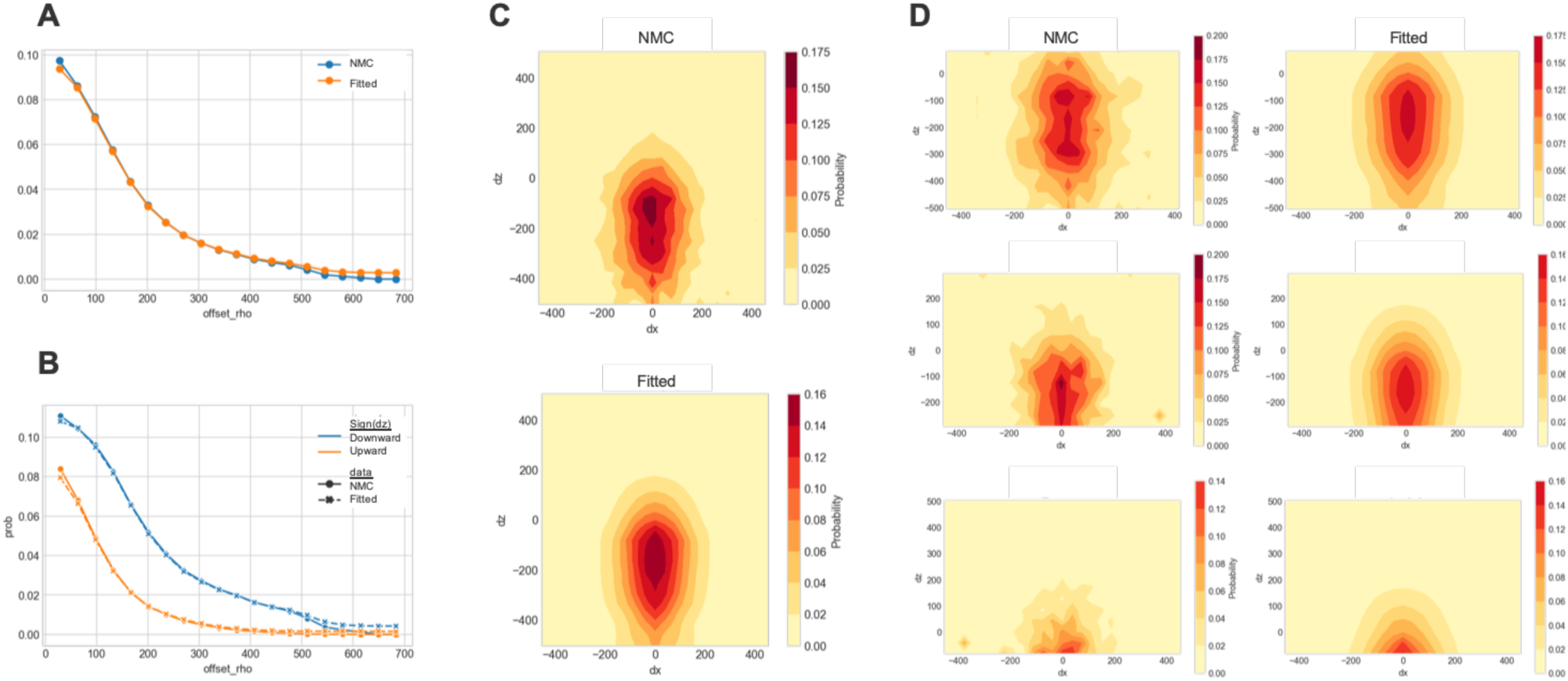
Comparison of empirical and fitted connection probabilities for L5-TTPCs. **a**, The dependence of connection probability on intersomatic distance as observed in the NMC (blue) and as fitted by the 2^nd^-order distance-dependent model (orange). **b**, The distinct connection probability of the downward (blue) versus upward (orange) connections as observed in the NMC (circle) and as fitted by the 3^rd^-order bipolar distance-dependent model (cross). **c**, The offset dependence of connection probability as observed in the NMC (top) and as fitted by the 4^th^-order offset dependent model. **d**, Same as in **c**, but for position dependence of the connection probability as fitted by the 5^th^-order position dependent model.

**Figure S2.**
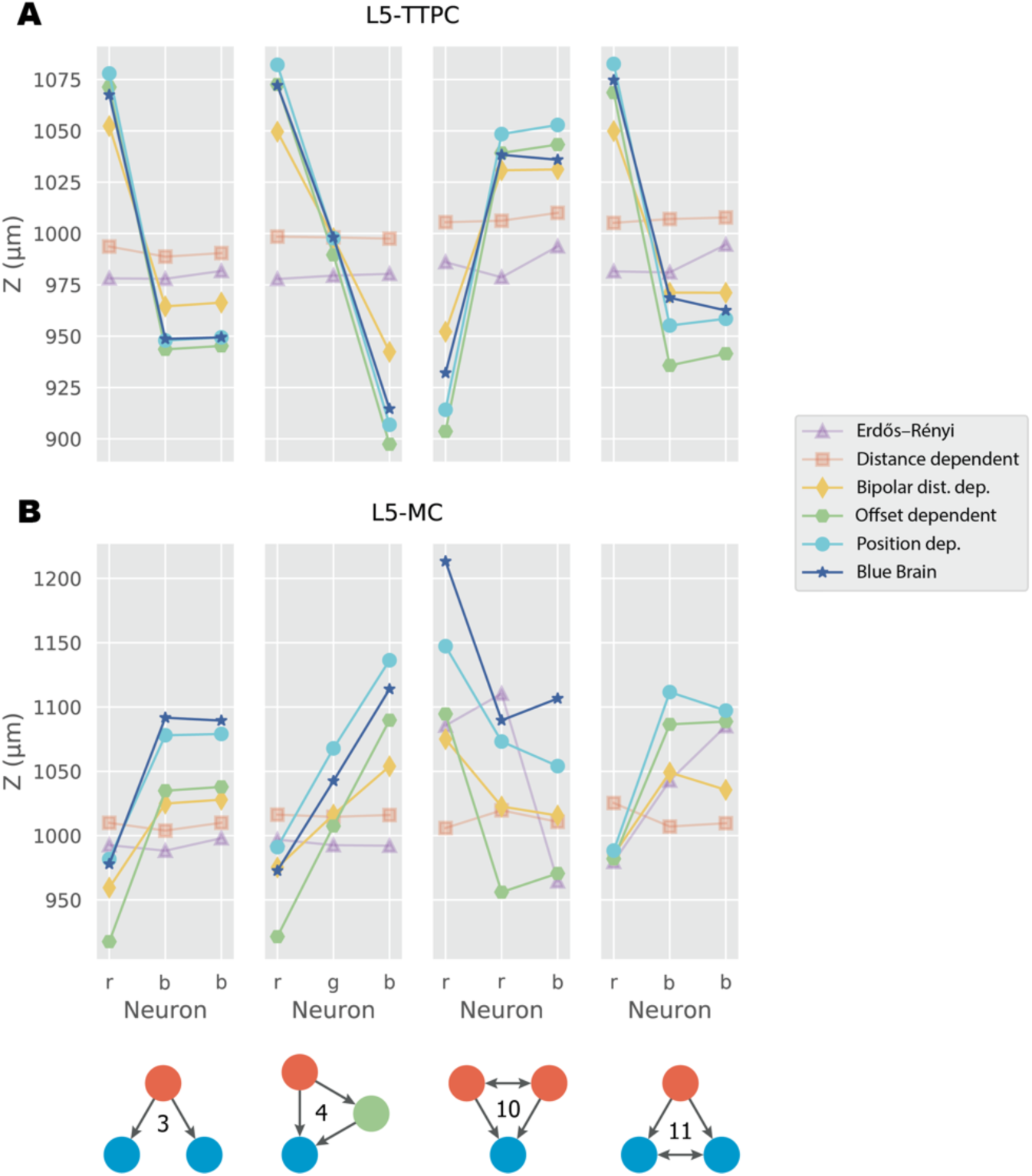
Spatial embedding of triplet motifs (#3, #4, #10 and #11) as predicted by the series of the geometric models. For **a**, L5-TTPCs and **b**, L5-MCs. For each model, an instance of one network was generated, on which the spatial alignment of the cells involved in forming the specific triplet was measured. Note that the anisotropic models (beyond the 2^nd^-order distance-dependent model) predict the asymmetrical embedding found in the NMC model (blue) and in the slice (**Fig. 4**). For instance, the embedding of L5-TTPCs feedforward motif (#4) is such that the “source” neuron (red node) is located above the other two cells whereas the “sink” neuron (blue node) is the lowest.

## Notes

### Competing Interest Statement

The authors have declared no competing interest.

### Summary of Updates

Text revisions.

